# Gene editing in “cell villages” enables exploring disease-relevant mutations in many genetic backgrounds

**DOI:** 10.1101/2025.11.08.687374

**Authors:** Rachel A. Battaglia, Sonia Bolshakova, Ilinca Mazureac, Dhara Liyanage, Noah Pettinari, Autumn Johnson, Ethan Crouse, Sartaj Habib, Isabel Flessas, Ajay Nadig, Derek Hawes, Matthew Tegtmeyer, Caroline Becker, Sulagna Ghosh, Giulio Genovese, Marina Hogan, Adrianna Maglieri, Lindy E. Barrett, Laurence Daheron, Steven A. McCarroll, Ralda Nehme

## Abstract

Understanding how individual genetic backgrounds shape the effects of disease-associated mutations is central to elucidating the biology of complex psychiatric disorders. We developed a scalable ‘village editing’ strategy that enables simultaneous genome editing across multiple induced pluripotent stem cell (iPSC) lines, allowing systematic assessment of how polygenic context modulates the impact of specific mutations. Using pooled CRISPR editing in 15 iPSC lines spanning a range of schizophrenia (SCZ) polygenic risk scores, we generated homozygous and heterozygous knockouts in two known SCZ-associated genes: *LRP1*, involved in cholesterol import, and *NRXN1*, a presynaptic adhesion molecule. By mixing all lines prior to editing and de-multiplexing them afterward, we efficiently produced multi-donor knockout neurons at scale. Transcriptomic profiling revealed that *LRP1* and *NRXN1* loss produce both shared and donor-specific effects on neuronal gene expression, with variable perturbation of neurotransmitter transport and cholesterol biosynthesis pathways across genetic backgrounds. These results demonstrate that village editing enables systematic dissection of gene-background interactions in human neurons, offering a powerful framework for studying the polygenic architecture of psychiatric disease.

## Introduction

Human disorders often exhibit complex genetic architecture that involves the contribution of a multitude of common risk factors. Because of this polygenic nature of many human diseases, the effect of one mutation can vary across many individuals. Identification and interpretation of genetic risk factors as well as understanding how they interact to manifest disease is one of the most significant challenges in the field of human genetic research. The intricate genetic landscape of human disease is exemplified by schizophrenia (SCZ), a debilitating and chronic psychiatric disorder that deeply impacts affected individuals and their families as well as society. Because of the high genetic heritability of SCZ^1^, large genetic studies have sought to elucidate the underlying factors driving the disease^2,3^. However, connecting genetic variants to molecular functions remains an ongoing challenge due to the prevalence of non-coding variants as well as the polygenic nature of SCZ. This, in turn, has stalled the development of effective therapeutics. There is thus a need to develop sophisticated tools that take interindividual genetic variability into account to functionalize these rich genetic resources and identify novel pathogenic mechanisms.

While the field of iPSC gene editing has experienced many technological improvements over the past decade^4–6^, it remains time consuming and costly to generate knockout (KO) cells in several different iPSC lines. Here, to address these limitations and screen different genetic backgrounds, we developed village gene editing, a scalable, efficient method for generating mutations in many iPSC lines simultaneously. As a proof-of-concept, the effect of knockouts of two known SCZ-associated genes, *Neurexin 1* (*NRXN1*) and *Low Density Lipoprotein (LDL) Receptor Related Protein 1* (*LRP1*) was studied in multiple genetic backgrounds. The development of this village editing approach allows the consideration of common risk alleles that may influence the manifestation of *LRP1* and *NRXN1* mutations through the selection and editing of several male and female donor lines with low, neutral, or high polygenic risk score (PRS) for SCZ.

Astrocyte-neuron interactions have been newly implicated as a key pathway in SCZ and other psychiatric and neurodevelopmental disorders such as autism spectrum disorder^7–10^. In a human neuron-mouse astrocyte co-culture system^11^, synaptic gene-expression programs were enhanced in neurons co-cultured in physical contact with astrocytes^8^. Additionally, co-culture resulted in the induction of genes associated with cell adhesion and cholesterol biosynthesis in astrocytes–pathways that were enriched for genes harboring variants associated with SCZ^8^. In human brain tissue (dorsolateral prefrontal cortex) sampled from 190 people, these same gene-expression pathways appear to fluctuate together across people^7^, with astrocytic expression of genes involved in cholesterol biosynthesis and synaptic cell adhesion fluctuating together with neuronal expression of genes encoding synaptic components. These are part of a larger coordinated gene expression program, the Synaptic Neuron and Astrocyte Program (SNAP)^7^. Expression of SNAP declines with advancing age and in persons with SCZ^7^. The genes recruited by SNAP in astrocytes are highly enriched for genes implicated by common and rare variants in genetic studies of SCZ^7^.

As a proof-of-concept study, we therefore assessed how genetic background affects the possible functions of synaptic cell adhesion and cholesterol biosynthesis pathways in SCZ. As a genetic perturbation of synaptic cell adhesion, we selected *NRXN1*, which encodes a presynaptic cell adhesion protein. NRXN1 contains a cytoplasmic PDZ protein binding domain and extracellular Laminin-a, Neurexin, and sex hormone-binding globulin (LNS) domains, enabling the formation of intra- and inter-neuronal complexes involved in synaptic organization and neurotransmitter release^12,13^. Recurring deletions of *NRXN1* exons greatly increase risk for SCZ as well as autism spectrum disorder (ASD) and many other neurodevelopmental disorders^14–16^, demonstrating pronounced phenotypic variability that could in principle result from differences in genetic background. Additionally, single nucleotide polymorphisms in *NRXN1* have been shown to affect individuals’ responsiveness to clozapine^17,18^, a second generation antipsychotic commonly used in cases of treatment resistant SCZ^19^, indicating that *NRXN1* could be a translationally relevant target. As a genetic perturbation of the astrocyte-to-neuron cholesterol shuttle, we chose *LRP1*, which is an endocytic receptor that binds to Apolipoprotein E and is one of the major receptors responsible for cholesterol import into neurons^20,21^. Risk of SCZ associates with multiple (linkage disequilibrium-independent) SNPs at the *LRP1* locus^3^. In addition to cholesterol, LRP1 has also been shown to bind and promote uptake of tau and a-synuclein^22,23^, and a single nucleotide polymorphism (SNP) in *LRP1* along with protein truncating variants have recently been linked to SCZ^24–26^.

Heterozygous and homozygous KOs in *NRXN1* and *LRP1* were generated in iPSCs from 15 donors, and subsequently, these cells were differentiated to neurons and analyzed for transcriptional alterations. *NRXN1* and *LRP1* KO in neurons disrupted genes related to neurotransmitter release and exocytosis. The effect of each KO on transcriptomic changes varied greatly by donor, including the baseline expression of these pathways, highlighting the importance of examining multiple donor cell lines and providing a framework to understand the variable effect of the same mutation across different individuals. This work introduces village gene editing as a method to rigorously address a broad range of biological questions while considering the influence of human genetic background and develops new resources to explore the role of *NRXN1* and *LRP1* in human cells.

## Methods

### Human induced pluripotent stem cell (hiPSC) cohort and derivation and culture

The cohort of 18 donor hiPSC lines used in this study were either obtained from the California Institute of Regenerative Medicine (CIRM) hPSC Repository or reprogrammed at the Harvard iPS core from PBMCs collected by the laboratory of Dr. Deborah Levy at McLean Hospital (see Supplementary Table 1 for detailed information of cell line provenance and metadata). All McLean Levy (ML) iPSCs have been deposited in, and are available from, the NIMH Repository and Genomics Resource (NRGR). All relevant ethical guidelines have been followed, and any necessary IRB and/or ethics committee approvals have been obtained. Prior to gene editing, iPSCs were cultured and maintained in mTeSR1 media (STEMCELL Technologies, 85850) on Geltrex-coated (Thermo Fisher, A1413201) standard 6-well tissue culture plates (VWR, 62406-161). iPSCs were passaged with Accutase (STEMCELL Technologies, 07920). During gene editing and expansion, cells were cultured in mTeSR Plus media (STEMCELL Technologies, 100-0276) on Cultrux-coated (Trevigen, 3434-005-02) standard tissue culture plates and passaged using Accutase or Gentle Cell Dissociation Reagent (STEMCELL Technologies, 07174). All cultures were kept at 37°C, 5% CO2.

### Cell village assembly and gene editing

iPSCs from each donor were maintained as separate cultures for a week. Cells were dissociated with Accutase (STEMCELL Technologies, 07920) and mixed in equal proportions (1 x 10^6^ cells per donor). 5 x 10^5^ cells were collected for WGS sequencing with the NovaSeq 6000 system (Illumina) to determine donor representation (Fig. 1D) using the Census-seq algorithm^27^.

**Fig. 1.**
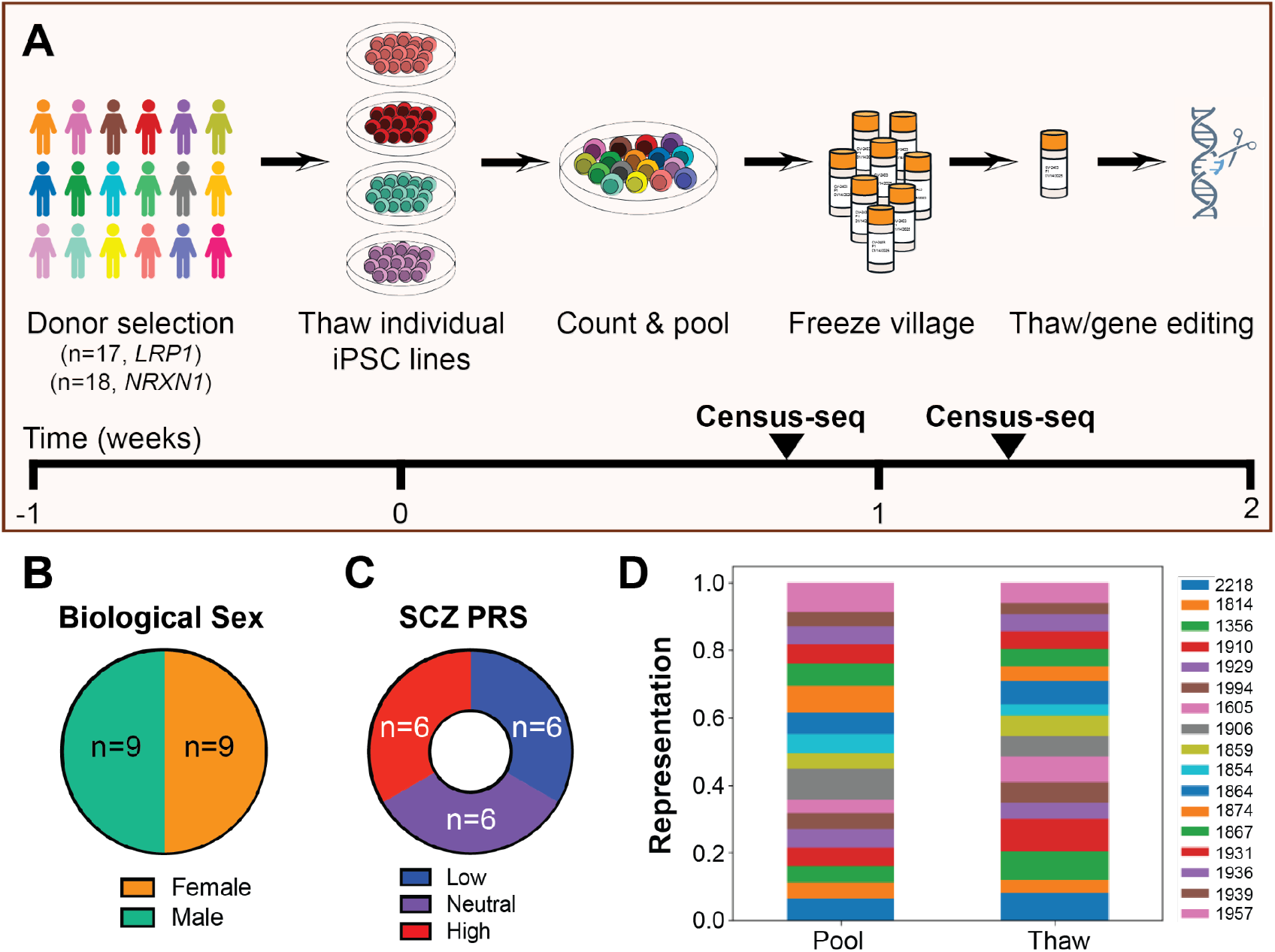
Design of cell villages for gene editing. **A** Schematic of cell village assembly and timeline. **B, C** Summary of donor information used to select cell lines for cell villages, including biological sex **B** and SCZ polygenic risk scores (PRS) **C**. The full donor information is available in Supplementary Table 1. **D** Confirmation of even donor representation in the cell village upon pooling and post-thaw by Census-seq^27^. The legend indicates which donors are represented by each color.

Two gRNAs were designed to target each gene (Supplementary Table 2), and gRNA cutting efficiency and electroporation conditions were optimized (Supplementary Fig. 1). Each gRNA (12.5 pmol) was assembled into an RNA-protein (RNP) complex with TrueCut Cas9 V2 (6.25 pmol) and electroporated into 1 x 10^6^ cells from the cell village with the Neon Electroporation system (pulse width: 30ms, 1 pulse). Electroporation was performed 2-3 times sequentially, and the cells pooled, for each of three voltages: 900V/100V/1100V for *NRXN1* and 900V/1100V/1300V for *LRP1*. To assess gRNA cutting efficiency, the target region was amplified by PCR from DNA isolated from pooled cells, and the amplicon was analyzed using either TIDE-coupled Sanger Sequencing with Next Generation Sequencing (*NRXN1*) or the Synthego Inference of CRISPR Edits (ICE) tool (*LRP1*). The best pools of cells, based on efficiency and viability (Supplementary Table 2, Supplementary Fig. 1, asterisks), were plated at low density to isolate clones. Colonies were picked and maintained for genotyping and cell expansion. Sanger sequencing and ICE analysis was used to determine the *NRXN1* and *LRP1* genotypes in order to select clones for expansion, and samples with insufficient viability, indels without frameshifts, or mixed genotypes were eliminated. The number of edited clones identified for each target are summarized in Supplementary Table 3 and Fig. 2C. Next, unedited clones as well as heterozygous and homozygous KOs were submitted for Illumina SNP Global Screening Array (GSA) for donor reidentification, and donor identities were validated by DNA fingerprinting. A summary of the edited clones identified for each donor is available in Supplementary Table 3. One clone for each genotype per donor was tested for mycoplasma, re-genotyped, and expanded for future experiments.

**Fig. 2.**
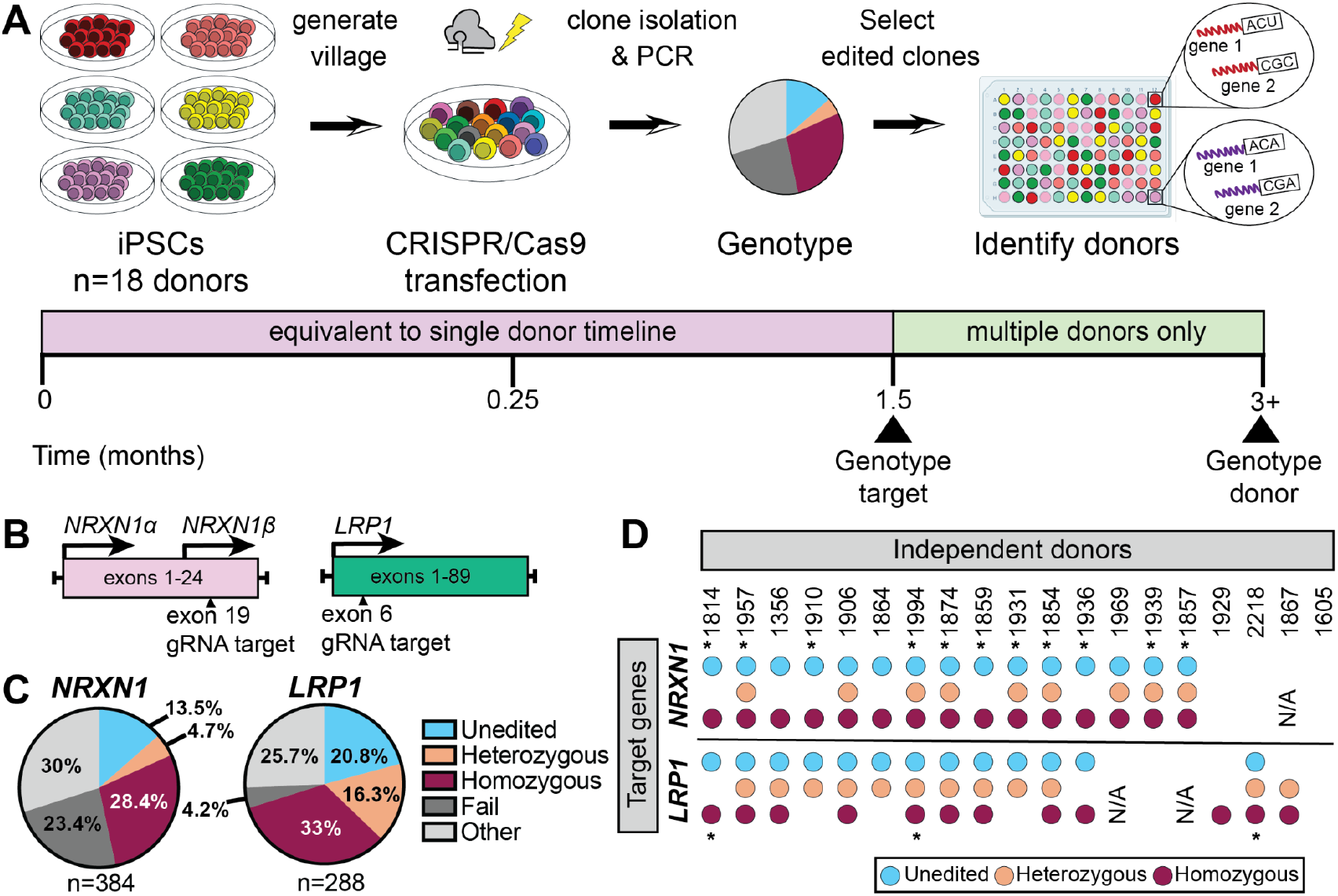
Cell village gene editing enables scalable production of *NRXN1* and *LRP1* gene knockouts in iPSCs. **A** Workflow for cell village gene editing procedure. **B** Schematic of *NRXN1* and *LRP1* genes and exons targeted with CRISPR-Cas9. The gRNA target cut site is indicated by an arrowhead for each gene. **C** Summary of gene editing efficiency results from the village editing screen. Unedited as well as heterozygous and homozygous *NRXN1* and *LRP1* KO cell lines were obtained. Fail indicates failed sequences and other signifies unwanted events such as indels without frameshifts or mixed clones. **D** Summary of genotypes obtained across different donor cell lines. Successfully acquired genotypes are indicated by the presence of blue (unedited), yellow (heterozygous KO) and red (homozygous KO) circles. Asterisks (*) indicate cell lines that were used for Fig. 3 and 4. N/A indicates a cell line that was not included in the original village that went through the editing process.

### Neuronal differentiation

For each cell line, the doxycycline-inducible *NGN2* construct was integrated into the *AASVI* safe harbor locus through NEON electroporative transfection (Thermo Fisher, MPK10016)^11,80^. hiPSCs were differentiated into cortical glutamatergic neurons using previously described *NGN2*-based method, which combines *NGN2* over-expression with small-molecule patterning to create a highly homogenous population of post-mitotic excitatory neurons^11^. At 4 days post induction, cells were co-cultured with mouse glia to promote neuronal maturation and synaptic connectivity^81,82^. All experiments were performed at 28 days post induction unless otherwise noted.

### Genotyping and PRS calculation

Whole-genome sequencing (WGS) data was collected for each donor used in the experiment. The WGS variant call format (VCF) was filtered according to the following parameters and thresholds: filtering variants with variant quality score recalibration (VQSR) not equal to PASS; allelic depth < 10; genotype quality < 20; allele balance filtering of: ref site > 10% alt reads, alt site > 10% ref reads, het site < 25% ref reads or >75% ref reads; Hardy-Weinberg Equilibrium (HWE) error rate <1e-3; call rate < 0.5; filtering variants in low complexity regions; filtering variants located in segmental duplication regions; removing variants in common exclusion sites in the GRCh38 reference genome. Samples in the VCF were filtered out if they had a sample call rate < 97% or sample median depth < 10.

After the village gene editing, concordance analysis was performed by sequencing the edited lines via SNP array (Illumina GSA). After filtering of duplicate samples, the GSA VCF was converted to PLINK format using the bcftools liftover plugin (for liftover from GRCh37 to GRCh38) and the vcf2plink.py script from the gwaspipeline GitHub repository. Concordance for each donor was validated using bcftools gtcheck (v.1.20) against the WGS sequencing data to identify putative sample swaps, duplicates, or mixed samples between cell lines from different donors. All sequenced cell lines were confirmed concordant with their donor of origin before being used in experiments.

Polygenic risk scores (PRS) for cell lines were calculated from genotypes generated by the GSA microarray. Genotypes at untyped loci were imputed using the 1000 Genomes Project haplotype reference panel^83^. The scores were calculated using summary statistics from the largest schizophrenia cohort analyzed at the time of this writing^3,84^. The PRS were calculated using linear regression betas for approximately 7,400 variants that achieved a p-value cut-off threshold of 0.01 in the discovery cohort, after variants were pruned to account for linkage disequilibrium patterns. Scores were further adjusted for sex and for the 10 first principal components computed after merging cell lines genotypes together with genotypes from the 1000 Genomes Project^83^ samples using an established protocol (https://github.com/freeseek/kgp2anc). Scores less than −1 or greater than 1 were classified as low and high PRS, respectively, whereas scores falling between −0.5. and 0.5 were considered neutral PRS.

### RNA-sequencing and processing

RNA was collected from D28 neuron-astrocyte co-cultures using the Qiagen RNeasy kit (Qiagen, #74106) and underwent bulk RNA sequencing at the Broad Genomics Platform according to the Smartseq2 workflow (cite PMID: 24385147). Raw paired-end RNA-seq FASTQs were processed using a custom bulk RNA-seq workflow. Adapter sequences were marked using Picard MarkIlluminaAdapters (v3.1.0), and aligned to the ENSEMBL GRCh38/GRCm38 combined reference genome and GENCODE GTF annotation version 19 using STAR (v2.5.2a). The output BAM files were then demultiplexed with Picard’s MergeBamAlignment. After marking PCR duplicates, reads in each transcriptome-aligned BAM file were quantified using RSEM’s (v1.3.3) rsem-calculate-expression. Estimated counts were then used to create two DGE matrices for each batch: one for human neuronal reads, and one for mouse glia reads. DGEs were then concatenated within each species across sequencing batches.

### Differential expression analysis

Differential gene expression (DGE) analysis was performed on bulk RNA-seq data using DESeq2 (v1.32.0) in R. Raw gene counts were pre-filtered with edgeR’s filterByExpr (v3.36.0) to remove lowly expressed genes (≤10 counts in any sample or ≤15 total counts across all libraries). Known sex-linked genes (DBY, SMCY, UTY, RPS4Y, USP9Y, XIST^85^) were excluded to prevent sex-related confounding. Counts were normalized using DESeq2’s median-of-ratios method. Due to limited donor availability across all three genotypes, LRP1 DGE analysis focused on pairwise comparisons between *LRP1* −/− and *LRP1* +/+. A linear model was constructed with genotype as the sole covariate (∼ 0 + target_gt). Variance-stabilizing transformation (VST) was applied for principal component analysis (PCA), and differential testing was conducted using Wald statistics. For NRXN1, two complementary approaches were implemented – pairwise contrasts and dosage-based modeling. For pairwise assessments, DE was assessed using a model analogous to that used for LRP1 (∼ 0 + target_gt), comparing both *NRXN1* −/− vs. *NRXN1* +/+ and *NRXN1* +/-vs. *NRXN1* +/+. For dosage-based modeling, a subset of five donors harboring all three NRXN1 genotypes was selected to investigate genotype dosage effects. A numeric dosage variable was encoded (0 = +/+, 1 = +/-, 2 = −/−), and a linear model of the form ∼genotype_dosage was used in DESeq2 to identify genes exhibiting monotonic expression changes in response to gene dosage. Donor was included as a covariate when multiple donors were analyzed together, but it was omitted when analyzed separately. Across all analyses, p-values were adjusted using the Benjamini–Hochberg procedure to control the false discovery rate (FDR) at 5%.

### Gene set enrichment analysis

Gene set enrichment analysis (GSEA)^30^ was performed on the DGE results obtained from DESeq2. For each differential expression contrast, genes were ranked by their log2 fold change values. The fgsea package (v1.18.0) in R was used for the analysis. Pathways with a gene count between 15 and 500 were considered (minSize = 15, maxSize = 500). Enriched gene sets were then adjusted for multiple testing using the Benjamini-Hochberg correction to control the False Discovery Rate (FDR) at 5%. Results were visualized by plotting the Normalized Enrichment Score (NES) for each pathway, with point size indicating statistical significance (-log_10_[FDR]) and point color indicating NES directionality.

Gene sets were utilized from several collections: the Hallmark gene sets (v2024.1) and Canonical Pathways (v2024.1) from the Molecular Signatures Database (MSigDB). Additionally, a custom-compiled gene set was used, which integrated the C5 Gene Ontology collection (v7.4) ^86,87^ from the Molecular Signatures Database^88,89^, SynGO (release 20210225) biological process (BP) and cellular component (CC) gene lists, and a custom gene set comprising genes implicated in schizophrenia genetic studies of humans (including genes at 1-2 gene loci from the SCZ GWAS (PGC3)^3^ and genes containing rare coding variants from SCHEMA^2^; FDR < 0.05).

### Latent factor analysis

Latent factor (LF) analysis was performed using Probabilistic Estimation of Expression Residuals (PEER)^44^, using R v3.5. Donors with fewer than three replicates and samples with fewer than 25% reads mapping to coding regions were excluded. For each dataset, raw gene expression counts from neurons and astrocytes separately were first normalized using counts per million (CPM), followed by log transformation and rank-based inverse normal transformation (INT). The top 5000 most variable genes were selected separately from mouse and human matrices by computing per-gene variance across samples to reduce noise and focus on more biologically relevant results. These matrices were merged, resulting in a combined expression matrix containing approximately 7000 genes, reflecting partial overlap between cell types. PEER was run on the merged expression matrices using 15 LFs.

PEER factor loadings were visualized with heatmaps, stratifying samples by genotype for each cell type. Heatmaps were generated both in aggregate (averaged across all donors per genotype) to reveal global genotype-dependent shifts, and separately for each donor to examine inter-individual variation. For individual PEER factors, genotype-wise comparisons of factor distributions between genotypes were performed using pairwise Wilcoxon rank-sum tests implemented and visualized with the geom_signif function from the ggsignif R package^90^. GSEA was performed on the PEER weight matrix using the fgsea package in R(v1.18.0). Enrichment analysis was performed independently for each latent factor, enabling identification of pathways driving variance captured by PEER components.

### qPCR analysis

Cell Lysates were prepared from neurons 14 days post induction (NRXN1) or iPSCs (LRP1) cultured on standard 6-well tissue culture plates (VWR, 62406-161). RNA was isolated using the RNeasy Kit (Qiagen, #74106) and converted to cDNA using the iScript cDNA Synthesis Kit (Bio-Rad, #1708890). The samples were prepared for qRT-PCR using the iQ SYBR Green Master Mix (Bio-Rad, #1708880). The primers used to detect gene expression in this study are listed in Supplementary Table 2.

## Results

### Village editing: a scalable method to generate knockout iPSC lines

To examine the functions of *NRXN1* and *LRP1* in different genetic backgrounds, 18 cell lines were selected from male and female donors with low, neutral, and high polygenic risk score (PRS) for SCZ (Fig. 1A-C, Supplementary Table 1, Methods). Next, “cell villages” were assembled by pooling equal numbers of iPSCs from each donor together and freezing multiple vials for future experiments (Fig. 1A). Even representation of donors within the cell village was confirmed by Census-seq^27^ pre- and post-thaw (Fig. 1D).

Following cell village assembly, gene editing was conducted using CRISPR/Cas9 to generate knockouts (KOs) in *NRXN1* and *LRP1* (Fig. 2A). Two guide RNAs (gRNAs) were designed to target each gene (Fig. 2B, Supplementary Table 2) and were tested for cutting efficiency (Supplementary Fig. 1). When designing gRNAs for *NRXN1*, the earliest possible common exon was targeted in the *NRXN1-α* and *NRXN1-β* isoforms, a previously validated approach^28^ which genocopies a SCZ patient^29^ (Fig. 2B). In the case of *LRP1*, a validated gRNA targeting exon 6 was utilized^23^. For village editing, preassembled CRISPR/Cas9 ribonucleoprotein complexes were electroporated into cell villages, and single cells were isolated for screening (Fig. 2A). PCR followed by Sanger sequencing was performed at the target region to genotype each clone, and clones with unedited (+/+), heterozygous KO (+/-), and homozygous KO (-/-) genotypes were selected for further analysis. Clones with desired genotypes were expanded and sequenced via SNP global screening array (GSA, Illumina) for donor identification and to confirm genomic integrity. After selecting clones from each donor for further expansion and downstream experiments, the donor of origin for each clone was further validated by DNA fingerprinting followed by concordance analysis. The *NRXN1* and *LRP1* knockouts were validated by qRT-PCR (Supplementary Fig. 2). The timeline to generate cell lines (3+ months, Fig. 2A) depends on the turnaround time of the sequencing data.

This method yielded an overall high efficiency of gene editing. Screening revealed 33.1% *NRXN1* KOs (n=384 total clones screened; *NRXN1-/-* = 109 clones, *NRXN1+/-* = 18 clones) and 49.3% *LRP1* KOs (n=288 total clones screened; *LRP1-/-* = 95 clones, *LRP1+/-* = 47 clones) (Fig. 2D, Supplementary Table 3). While differences were observed in donor representation after gene editing, KOs were obtained in 15 out of 18 donors for *NRXN1* and *LRP1* (Fig. 2D, Supplementary Fig. 3, Supplementary Table 3). These results show that the cell village gene editing method can rapidly and effectively produce KOs in over 80% of the desired donor cell lines for two different target genes.

### Transcriptional alterations in *NRXN1* KO neurons reveal inter-cell line variability

To begin characterizing the effects of *NRXN1* KOs, unedited (*NRXN1 +/+*) as well as any available *NRXN1 +/-* and *NRXN1 −/−* iPSC lines were used from a subset of donors (n=12 *NRXN1 +/+*, n=6 *NRXN1 +/-*, n=11 *NRXN1 −/−*) to generate neurons (Fig. 3A, Fig. 2D asterisks). All experiments following the initial village editing experiment were performed in an arrayed, non-pooled format, in quadruplicates for each condition and each donor. First, neural progenitor cells (NPCs) were produced and co-cultured with mouse glia and differentiated to neurons using established protocols^11^. At 28 days post induction (DPI), the cells were lysed, and bulk RNA-sequencing was performed. After excluding samples that did not pass quality control metrics (see Methods), conditions with a minimum of three technical replicates were included. The human neuron-mouse glia co-culture allowed for separation of neuronal and glial transcripts by cross-species transcript mapping. As expected, total expression of *NRXN1* was reduced in the *NRXN1+/-* and *NRXN1-/-* lines, and there were no significant differences in *NRXN1* expression related to PRS or biological sex (Fig. 3B, Supplementary Fig. 4).

**Fig. 3.**
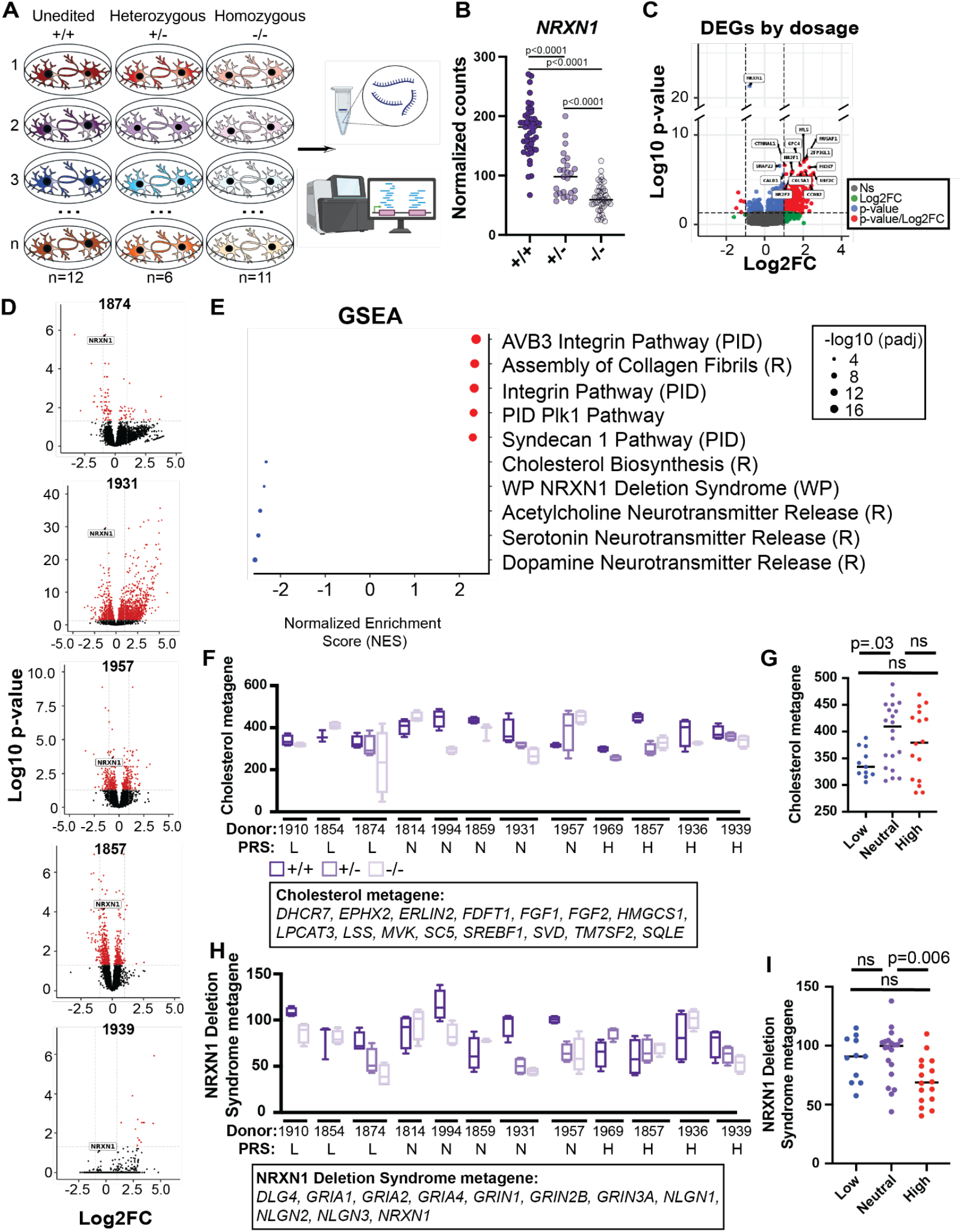
Characterization of transcriptional alterations in *NRXN1* KO neurons. **A** Experimental design for cross-species co-culture of *NRXN1 KO* human neurons with mouse glia. iPSCs from all available genotypes from each donor were plated in an arrayed format with mouse glia and were differentiated to neurons after which they were processed for RNA sequencing (n=12, 6, and 11 biological replicates for *NRXN1* +/+, +/-, and −/−, respectively, with 3-4 technical replicates each). **B** Confirmation of reduced *NRXN1* gene expression in the expected genotypes. **C, D** Volcano plots show differential gene expression analysis by *NRXN1* gene expression dosage in **C** all donors with every *NRXN1* genotype represented or **D** these same individual donors. **E** Gene set enrichment analysis (GSEA) of differentially expressed genes from **C** where upregulated pathways are red and downregulated pathways are blue. **F, H** Box plots showing metagene expression for cholesterol-(**F**) and NRXN1 Deletion Syndrome-(H) associated genes. **G, I** Dot plots showing cholesterol (**G**) or NRXN1 Deletion Syndrome (**I**) metagene expression in unedited samples from each PRS group. The p values are shown and were calculated by one-way ANOVA followed by Tukey’s multiple comparison test (**B**,**G**,**I**). For **C-E**, the Wald statistical test was applied, and enriched gene sets were then adjusted for multiple testing using the Benjamini-Hochberg correction to control the False Discovery Rate (FDR) at 5%.

Next, differential gene expression (DGE) analysis was performed comparing *NRXN1+/-* and *NRXN1-/-* to the unedited samples (Supplementary Fig. 5). Additionally, taking advantage of the cell lines for which all three genotypes were represented (n=5 cell lines), DGE was examined by *NRXN1* dosage (Fig. 3C, Methods). The dosage model enabled testing for genes whose expression changed in a monotonic, dose-dependent manner with increasing *NRXN1* copy loss. This method offered greater sensitivity than pairwise contrasts when detecting graded genotype effects. Interestingly, when differentially expressed genes (DEGs) were analyzed in individual cell lines, rather than as a group, the number and identity of DEGs varied greatly between cell lines (Fig. 3D, Supplementary Fig. 5, Supplementary Table 4), suggesting that genetic background may at least in part influence the transcriptional alterations in *NRNX1* deficient cells.

### Cholesterol and synaptic pathways are altered in *NRXN1* KOs and vary at baseline across cell lines

To identify biologically meaningful patterns among the gene expression changes, gene set enrichment analysis (GSEA) was performed (Fig. 3E, Supplementary Table 5)^30^. One of the downregulated pathways ranked most strongly by GSEA was “NRXN1 deletion syndrome” (GO: p-value = 4.67e-4, NES = −2.40), along with neurotransmitter release pathways, which are known to be disrupted in *NRXN1* knockout in mice and human cells^28,31^. The cholesterol biosynthesis pathway was among the top downregulated pathways, which was striking considering the importance of cholesterol-related pathways in neuron-astrocyte interactions and neuronal synapse and dendritic spine formation^7,32– 34^. Top GSEA pathways were examined across different cell lines by generating a metagene for each pathway, which is an average of the normalized gene expression of all genes in the pathway. Cholesterol-related genes exhibited not only variability in overall changes between unedited and KOs but also baseline differences across donors in unedited lines (Fig. 3F, 3G). When grouped by PRS, high SCZ PRS donors displayed higher expression of the cholesterol metagene at baseline (Fig. 3G). Strikingly, the NRXN1 deletion syndrome metagene, which includes many synapse-related genes (*GRIA1, GRIN2B, NLGN1*, etc.), was significantly reduced in high compared to neutral SCZ PRS donors at baseline (Fig. 3H, 3I). In fact, for the NRXN1 deletion syndrome metagene, the high SCZ PRS unedited lines had comparable expression levels to neutral and low SCZ PRS KO lines. These findings show that general gene expression changes (Fig. 3D) as well as disease-relevant pathways (Fig. 3F-I) can vary greatly at baseline in different cell lines, demonstrating the importance of using multiple donor cell lines with disease-relevant genetic backgrounds.

### KO of *LRP1* in neurons induces transcriptional alterations in neurons and in co-cultured astrocytes

Next, using the same differentiation strategy as above (Fig. 3A), *LRP1* KO lines (*LRP1* +/+ n=3, +/-n=2; −/− n=3 donors following RNA-sequencing quality control) were differentiated to neurons to explore gene expression changes related to *LRP1* depletion.

Knockdown of *LRP1* in neuronal progenitor cells (NPCs) using CRISPRi in a single iPSC line was shown to result in gene expression changes in the co-cultured glial cells two days after co-culture^32^. To extend these observations, the same cross-species co-culture was leveraged to examine gene expression changes in both more mature neurons (D28 post-induction) and glia. DGE analysis was performed comparing *LRP1* −/− to +/+ neurons as well as comparing wild-type mouse glia co-cultured with *LRP1 −/−* versus *LRP1* +/+ neurons (Supplementary Table 6). First, the differentially expressed genes were examined in neurons (Fig. 4A, Supplementary Fig. 6A, Supplementary Table 7). While DEGs varied across cell lines (Supplementary Fig. 6A), as with *NRXN1* KO lines, the number of significant DEGs was greater in the grouped analysis for *LRP1* KO lines (Fig. 4A). As expected, *LRP1* was the most downregulated gene in neurons (Log2FC = −2.67, padj = 8.46e-9), which was confirmed by examining normalized counts of *LRP1* gene expression (Fig. 4B). Likewise, several cholesterol-related genes were among the downregulated genes, including *LSS* (Log2FC = −1.52, padj = 3.73e-6), *DHCR24* (Log2FC = −1.35, padj = 2.34e-4), *MVK* (Log2FC = −1.01, padj = 8.84e-4), *HMGCR* (Log2FC = −0.88, padj = 1.81e-4), and *PMVK* (Log2FC = −0.43, padj = 0.035) (Fig. 4B).

**Fig. 4.**
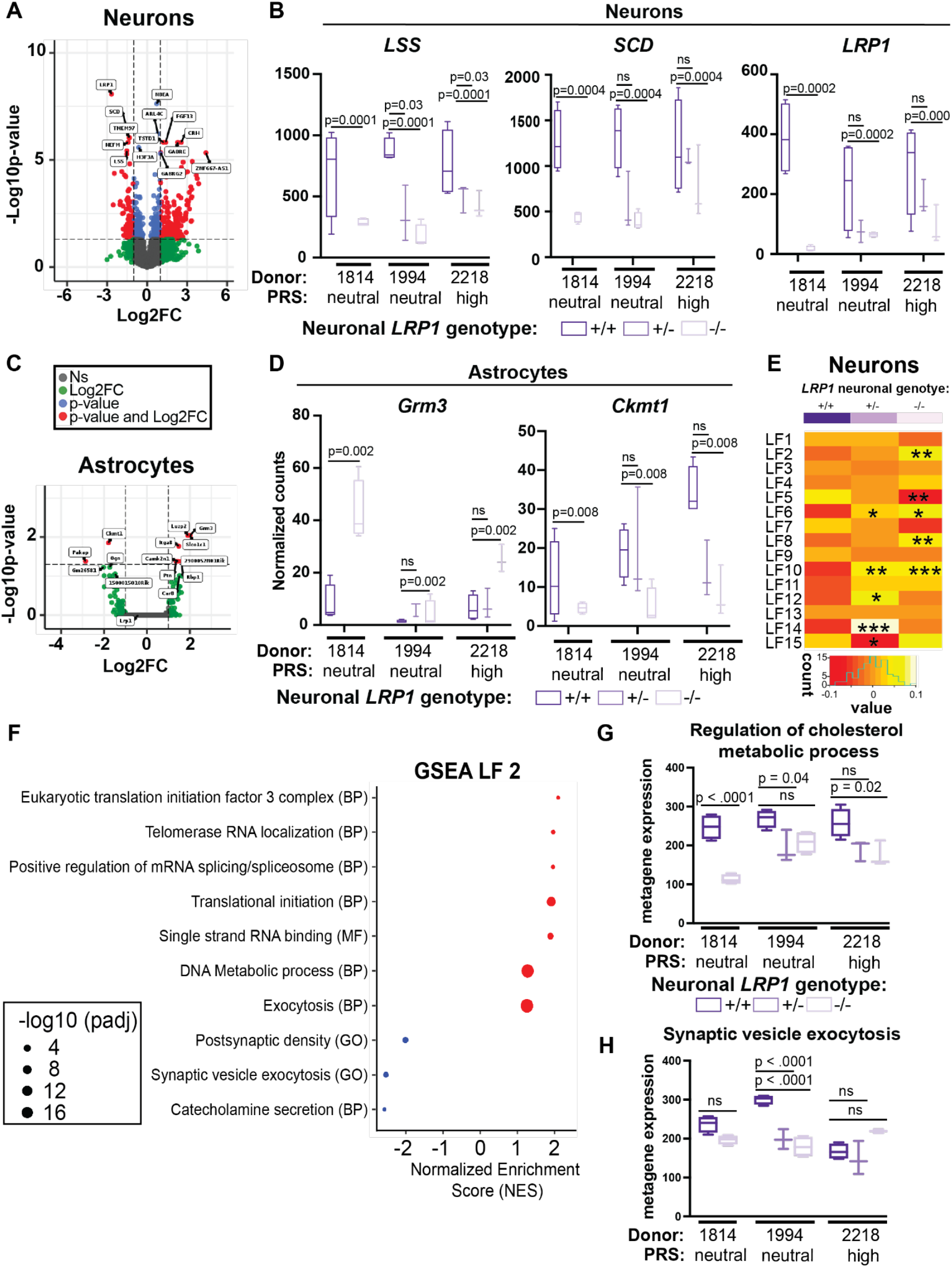
Depletion of synaptic vesicle trafficking genes in LRP1 KO neurons. **A, C** Volcano plots show DEGs in *LRP1* −/− neurons compared to unedited neurons (**A**) as well as astrocytes co-cultured with *LRP1* −/− neurons compared to astrocytes co-cultured with unedited neurons (**C**). **B, D** Box plots show normalized gene expression in neurons (**B**) or co-cultured glia (**D**) across donors and confirmation of reduced neuronal *LRP1* gene expression in the expected genotypes. **E** LF analysis of neuronal samples. **F** GSEA summary of LF2 where upregulated pathways are red and downregulated pathways are blue. **G, H** Box plots demonstrate cholesterol (**G**) and synaptic vesicle exocytosis (**H**) metagene expression in neuronal samples. For all panels, n=3 biological replicates (independent cell lines) for *LRP1* +/+ and *LRP1* −/−, and n=2 biological replicates for *LRP1* +/- with 3-4 technical replicates each. The p values were calculated by one-way ANOVA followed by Tukey’s multiple comparison test (**B**,**D**,**G**,**H**). For **A** and **C**, the Wald test was applied, and gene-level p-values were adjusted for multiple testing using the Benjamini-Hochberg correction to control the False Discovery Rate (FDR) at 5%. In panel E, \**p* < 0.05, \*\**p* < 0.01, and \*\*\**p* < 0.001 by Wilcoxon rank-sum test.

*LDLR*, another apolipoprotein endocytic receptor capable of cholesterol import^35^, was also downregulated in *LRP1* KO neurons (Log2FC = −1.59, padj = 5.98e-5). Alternatively, *LPL*, which encodes a lipoprotein lipase that hydrolyzes triglycerides^36^, was increased (Log2FC = 3.06, padj = 0.0012). These results are aligned with animal studies where neuronal *Lrp1* KO causes reduced brain cholesterol and triglycerides in mice^37^. Additionally, *TMEM97* was downregulated, which was notable because *TMEM97* has been associated with chronic pain^38^ and neuropathic pain injury-induced depression and anxiety^39^ (Log2FC = −1.34, padj = 9.01e-7).

Subsequently, differentially expressed genes were analyzed in mouse glia (Fig. 4C, Supplementary Fig. 6B, Supplementary Table 7). *Ckmt1*, encoding mitochondrial creatine kinase 1, was downregulated (log2FC = −1.79, padj = 0.014) (Fig. 4D). Deficiency in *CKMT1* has been shown to reduce mitochondrial function in human epithelial cells^40^. Mitochondrial-related gene expression was also elevated, which could indicate mitochondrial damage or dysfunction consistent with what has been observed upon cholesterol accumulation in astrocytes^41^. SNAP-related genes were also assessed by examining the top 500 gene loadings in astrocytes (SNAP-a) and neurons (SNAP-n) from Ling et al.^7^. In mouse glia, three SNAP-a genes were upregulated, including *Grm3* (log2FC = 2.01, padj = 0.009), *Ptn* (Log2FC = 1.35, padj = 0.04), and *Slco1c1* (Log2FC = 2.0, padj = 0.009). *GRM3* has been previously linked to SCZ^42,43^. In neurons, 16 SNAP-n genes were upregulated (*PTPN11, DDHD1, ELMO1, LPGAT1, SORL1, STS, SLC2A13, GPX3, MIR100HG, GTF2E2, LDLRAD4, CALU, FRMD6, PLS3, SERTM1*, and *CRH*) and 7 were downregulated (*BCL11A, LZTS1, GRM2, DNAL4, DUSP6, AFAP1*, and *ATAD2B*). Thus, human neuron (*LRP1* −/−) mouse glia (*LRP1* +/+) co-cultures show cholesterol-, SNAP- and disease-relevant changes in gene expression.

### *LRP1* KO neurons show decreased expression of lipid- and synaptic-related genes

To better understand how *LRP1* depletion affects astrocyte-neuron interactions, latent factor (LF) analysis was used to examine how gene expression changes fluctuate together in neurons (Fig. 4E) and astrocytes (Supplementary Fig. 7A) using 15 LFs^44^. GSEA was performed on the genes within each LF and found that unsurprisingly several included cholesterol- and lipid-related pathways (LF8, LF10, and LF11), which were consistently downregulated in neurons and upregulated in astrocytes (Supplementary Fig. 7, Supplementary Table 8). This suggested that cholesterol production and transport may be increased in astrocytes to compensate for the loss of neuronal LRP1, although the cholesterol metagene was not significantly changed in astrocytes (Supplementary Fig. 7B). Interestingly, LF2 and LF4 contained GSEA terms related to synaptic pathways, and LF2 was examined more closely because it was significantly increased in *LRP1* −/− neurons (Fig. 4F). LF2 had a negative enrichment for synaptic vesicle exocytosis genes, indicating a decrease in synaptic vesicle exocytosis gene expression. Additionally, LF2 decreased in astrocytes co-cultured with *LRP1 −/−* neurons, suggesting an increase in synaptic vesicle exocytosis genes (Supplementary Fig. 7). Further, LF4, which was positively enriched for many synapse-related GSEA terms (neurotransmitter secretion, synaptic membrane adhesion, dendrite morphogenesis) and SCZ-associated genes, was increased in astrocytes. The decrease in neuronally expressed synaptic genes and increase in astrocytic synapse-related genes extended findings of knockdown of *LRP1* in iPSC-NPCs by CRISPRi^32^. Metagene expression of the “regulation of cholesterol metabolic” pathway from LF8 and “synaptic vesicle exocytosis” pathway from LF2 were next examined across individual donor cell lines (Fig. 4F, 4G). While all cell lines exhibited reduced cholesterol metagene expression, only one cell line showed a significant decrease in synaptic vesicle exocytosis. This suggests that loss of *LRP1* may affect cell lines differently for a subset of transcriptional responses. Altogether, these results confirm that LRP1 plays an important role in neuronal cholesterol metabolism, and suggest its depletion impacts cholesterol metabolism as well as synaptic-related gene expression pathways in astrocytes and neurons.

## Discussion

We have described here a novel technique to rapidly and efficiently generate KO iPSCs from many donors simultaneously using the village gene editing method. To demonstrate the effectiveness of this method, two SCZ-related genes, *NRNX1* and *LRP1*, were targeted in 18 cell lines from donors with low, neutral, or high risk for SCZ, achieving high efficiency gene editing and donor coverage for both targets. This approach produced knockouts in over 80% of donor cell lines for both gene targets in mere months. All available iPSC lines were differentiated to neurons in co-culture with mouse glia. Gene expression changes varied greatly by genetic background, underlining the importance of including diverse genetic backgrounds in iPSC-based studies, especially those related to human diseases, such as SCZ, that are known to be polygenic.

Given the complex genetics of SCZ, it is unsurprising that individuals with gene variants associated with SCZ display phenotypic variability^45^. *NRXN1* clearly epitomizes this phenomenon, as copy number variants in this gene are associated with a broad array of disorders, including SCZ^14^, ASD^15,16^, attention deficit hyperactivity disorder^46^, intellectual disability^47^, Tourette syndrome^48^, epilepsy^49^, and depression^50^ and are also harbored by neurotypical individuals. These variable outcomes may be attributed to a combination of incomplete penetrance, gene pleiotropy, intersection with other genetic and environmental factors, and location of the mutation within *NRXN1*. Several groups have examined genotype-phenotype correlations for *NRXN1* and have found that the majority of exonic deletions occur at the 5’ end of the gene covering the *NRXN1-α* isoform, whereas deletions overlapping *NRXN1-β* are rarer^51–53^. However, individuals with SCZ and ASD have been identified harboring exonic deletions that affect both major isoforms, suggesting that both may play a role in disease pathogenesis^29,54^. These studies do not provide a clear, consistent explanation for the phenotypic variability. As suggested by Hu et al., one possible biological explanation could be that *NRXN1* isoforms can be differentially expressed in different brain regions or even synapses^51^. In fact, higher prefrontal cortex expression of *NRXN1-β* has been observed in SCZ and *NRXN1-α* in bipolar disease^55^. Interactions between NRXN1 and its post-synaptic ligands have also been shown to be isoform and splice variant dependent^12,13,56^. Isoform-specific interaction of NRNX1*α* with GABAergic-specific Calystenin-3^57,58^ and NRXN1*β* with GABAergic-specific Neuroligin 1^59,60^ has been observed. Likewise, the presence or absence of splice site 4, which can be present in both isoforms, is critical for many other interactions, adding further complexity ^61–64^. In fact, *NRXN1* is known to undergo alternative splicing resulting in thousands of variants, and recent evidence has implicated dysregulated alternative splicing of *NRXN1* in human disease^12,13,65^. The cellular tools presented here not only provide a means to understand the function of *NRXN1* but also create opportunities to study the role of specific *NRXN1* variants by overexpression into a *NRXN1α/NRXN1β* KO background. Importantly, future examination of natural variants of *NRXN1* and their altered splicing patterns will be a key complementary approach to understanding disease pathogenesis^66^.

An unexpected finding was the significant downregulation of the cholesterol biosynthesis pathway in *NRXN1* KO neurons. Brain cholesterol, which is predominantly provided by astrocytes, is essential for multiple neuronal processes, such as synaptogenesis, dendrite development, and axonal guidance^34,67^. A potential explanation for this downregulation may be altered neuronal demand for cholesterol. Given that gene expression results indicate *NRXN1* KO leads to impaired synaptic activity and synaptogenesis pathways in neurons, the need for cholesterol in neurons may decrease. Recently, a bidirectional signaling mechanism was described whereby neuronal cholesterol status directly influenced astrocyte gene expression and function^32^. Neuronal knockdown of *LRP1* not only downregulated neuronal synaptic genes but also induced a transcriptional response in astrocytes, including upregulation of *Nrxn1*^*32*^. In the current study, *NRXN1* dosage analysis identified altered cholesterol biosynthesis pathways in neurons, which, despite donor-to-donor variability, supports the idea of a neuron-astrocyte feedback loop. Cholesterol pathways also had decreased expression in *LRP1* KO neurons, in agreement with previous work showing reduced brain cholesterol and triglycerides in mice with conditional KO of *Lrp1* in neurons^37^. It has been shown that impaired cholesterol synthesis in astrocytes can lead to reduced dendrite growth from neurons^33,34,68^. While previous studies focused on rapid changes after 2 days of co-culture, these findings are extended to older, fully differentiated neurons. Changes in cholesterol metabolism in neurons and astrocytes, which were not detected in the younger cultures^32^, may occur on a longer timescale. This proof-of-concept experiment had a limited number of donors and found no major differences in cholesterol pathways across PRS categories but a slight reduction of synaptic vesicle exocytosis pathways at baseline in the high SCZ cell line (Fig. 4F,G). It will be important to examine this in more donors in order to understand the significance of these findings in the context of genetically stratified donor backgrounds. Our work provides key cellular resources that can be used to further understand the role of cholesterol and synaptic cell adhesion pathways in astrocyte and neuron functions using disease-relevant genetic backgrounds.

Lastly, we note that, while this method was applied here to generate knockouts that disrupt gene function, the strategy has broad applicability to different kinds of mutations and disease contexts. For instance, cell villages could be combined with base editing^72^ to generate point mutations, or homology-directed repair to generate larger knock-ins, and the use of iPSCs allows for generation of multiple cell types to study a wide range of human diseases.

Previous work using cell village systems underscored the challenge of maintaining stable donor representation over long periods in dividing cells such as iPSCs^26,69^. This is consistent with the findings that the composition of donor cell lines changed after gene editing (Fig. 2D, Supplementary Fig. 3) and motivated us to perform post-editing clonal isolation and expansion in the current study. Future optimization of the workflow (for instance, via optimization of electroporation conditions to increase viability) may enable differentiation directly in the pooled village format to further increase the scalability. Some donors lacked either the +/- or −/− KO genotype, which limits the feasibility of examining dosage-dependent effects in these specific backgrounds. Interestingly, the +/-genotype was underrepresented in the *NRXN1* targeted lines but not across *LRP1* targeted lines (Fig. 2D) This is a documented phenomenon, and the discrepancy is likely related to differences of gRNA cutting efficiency or the chromatin landscape at the *NRXN1* and *LRP1* loci^70^. Distance of gRNA sequence from the intended target can also contribute to rates of heterozygosity versus homozygosity in homology-directed repair^71^. These obstacles can be surpassed by testing different electroporation voltages or gRNAs to obtain heterozygotes. Lastly, this study had limited power to quantitatively assess the impact of individual genetic backgrounds or factors (e.g., sex or PRS category), owing to the small number of samples within each subgroup. The application of rigorous quality control measures at multiple stages of the workflow further reduced the number of lines available for analysis, particularly within the *LRP1* cohort. Future work will benefit from scaling both the initial size of the villages and the number of edited lines screened using cost-effective approaches, thereby enabling quantitative evaluation of how distinct genetic background factors contribute to specific phenotypes.

Incorporating genetic diversity in stem cell research is a uniquely promising approach to accelerate scientific discovery, and expand therapeutic strategies^73–76^. Our motivation to uncover novel pathways related to the pathogenesis of SCZ led us to include iPSCs from donors with low, neutral, or high PRS for SCZ. Because PRS calculations are affected by ancestry^76^, this study was limited to individuals of European ancestry for which the highest sample size, and thus the most accurate PRS calculations, were available^3,77,78^. As ancestry representation in GWAS and repositories improve, including cell lines from donors of various ancestries will be important.

In summary, we developed cell village gene editing, a new method that can be used to rapidly generate several KO iPSC lines simultaneously. Using this method, *NRXN1* and *LRP1* KOs were generated in 15 iPSC lines, and available lines were differentiated to neurons, which show gene expression differences at baseline across different donor cell lines. This method has potential to be used in a broad range of iPSC models of polygenic human disease to study interindividual differences in KOs as well as point mutations and other knock-in mutations. These findings underline the importance of including many different donors in iPSC-based studies in order to perform robust studies and provide an accessible roadmap for accomplishing this in a time- and cost-effective manner.

## Acknowledgments

This work was supported by NIH grants R01MH128366 (R.N.), U01MH115727 (R.N., S.A.M.), and K00AG068523 (R.B.), the Broad Institute Next Generation Award (R.N.) and the Stanley Center for Psychiatric Research Gift (R.N.). We thank Kris Dickson for reviewing and editing the manuscript, and Sovann Woodin for technical assistance with genotyping.

## Author contributions

R.N., R.B., S.B., I.M., and D.L. designed experiments and wrote and edited the manuscript with input from all authors. I.M. and D.L., with some help from M.T., conducted the neuronal differentiation experiments and collected data. S.B., N.P., and R.B. analyzed transcriptional data. S.G. and G.G. performed polygenic risk score calculations. L.D. oversaw the gene editing process and C.B. conducted the gene editing. R.B., I.M., D.L., A.J., E.C., S.H., I.F., and D.H. generated, maintained and genotyped iPSC lines. M.H. and A.M. managed and coordinated responsibility for the research experiment planning and execution. R.N. and S.M. conceived the project and R.N. supervised the work and analysis.

## Availability of Data and Materials

Requests for further information should be directed to the lead contact, Ralda Nehme.

