## Supplemental Figure 1 for "Gene editing in “cell villages” enables exploring disease-relevant mutations in many genetic backgrounds"

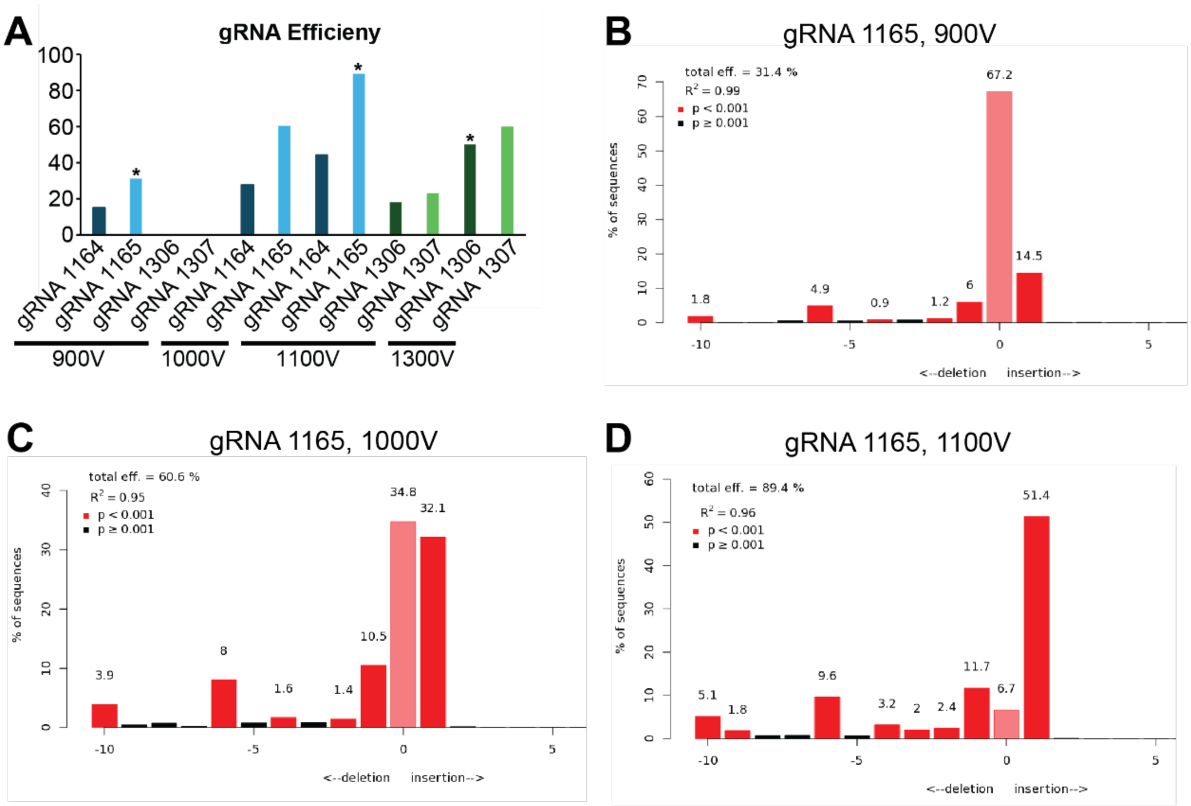


**Supplementary Fig. 1 | Optimization of electroporation of gRNAs, related to Fig. 2.** **A** Summary of the percent gRNA cutting efficiency measured by either TIDE (*NRXN1*, blue) or ICE analysis (*LRP1*, green), which is listed in Supplementary Table 3. Asterisks indicate conditions utilized for cell village CRISPR editing, not p values. **B-D** Determination of *NRXN1* 1165 gRNA cutting efficiency by TIDE-coupled Sanger Sequencing with Next Generation Sequencing[79]. The percentage of insertions and deletions detected in DNA sequenced from pooled iPSCs edited with the indicated gRNA and electroporation conditions. The overall efficiency is indicated at the top of each graph in parentheses.
