## Supplemental Figure 2 for "Gene editing in “cell villages” enables exploring disease-relevant mutations in many genetic backgrounds"

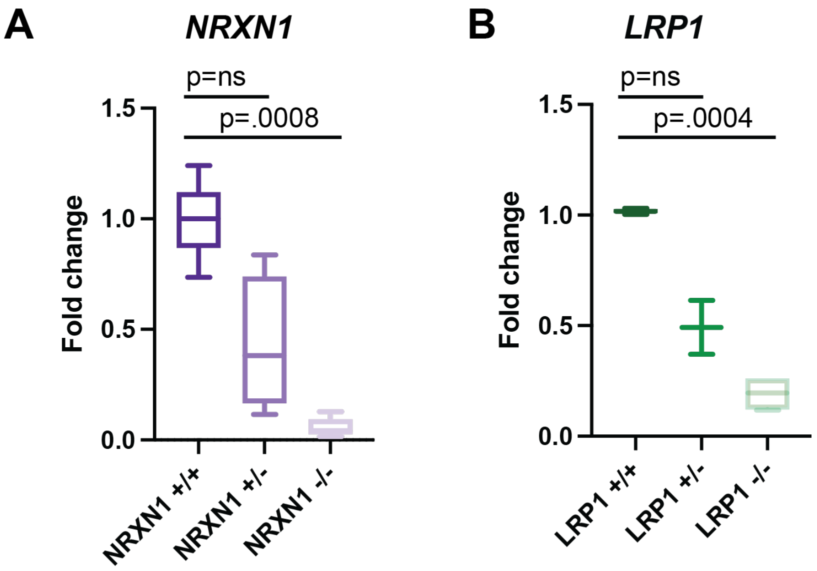


**Supplementary Fig. 2 | Validation of *NRXN1* and *LRP1* knockout cell lines, related to Fig. 2. A, B** qRT-PCR shows that *NRXN1* (**A**) and *LRP1* (**B**) gene expression are decreased in the KO genotypes. The p values are shown and were calculated by one-way ANOVA followed by Tukey's multiple comparison test (n = 5 biological replicates, 3 technical replicates).
