## Supplemental Figure 3 for "Gene editing in “cell villages” enables exploring disease-relevant mutations in many genetic backgrounds"

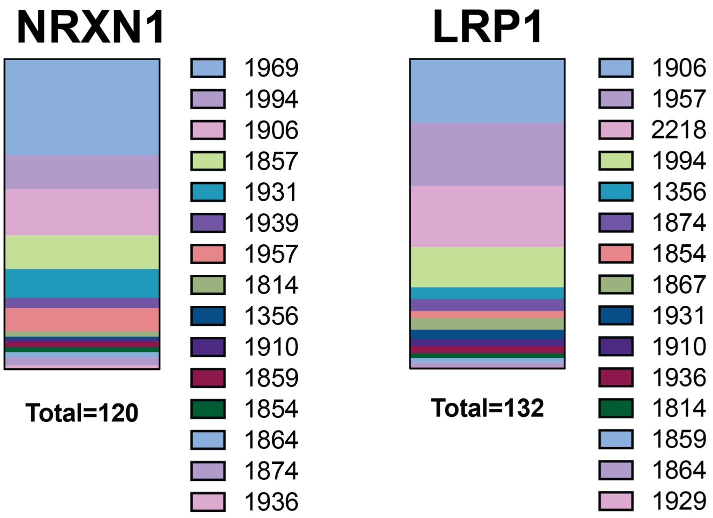


**Supplementary Fig. 3 | Donor representation after gene editing, related to Fig. 2.** Bar graphs show the proportion of donors identified in post-editing screening clones (n=120 clones screened, *NRXN1*; n=132 clones screened, *LRP1*). Each sample included represents a clone that was genotyped and contained the desired genotype (unedited, heterozygous KO, or homozygous KO).
