## Supplemental Figure 4 for "Gene editing in “cell villages” enables exploring disease-relevant mutations in many genetic backgrounds"

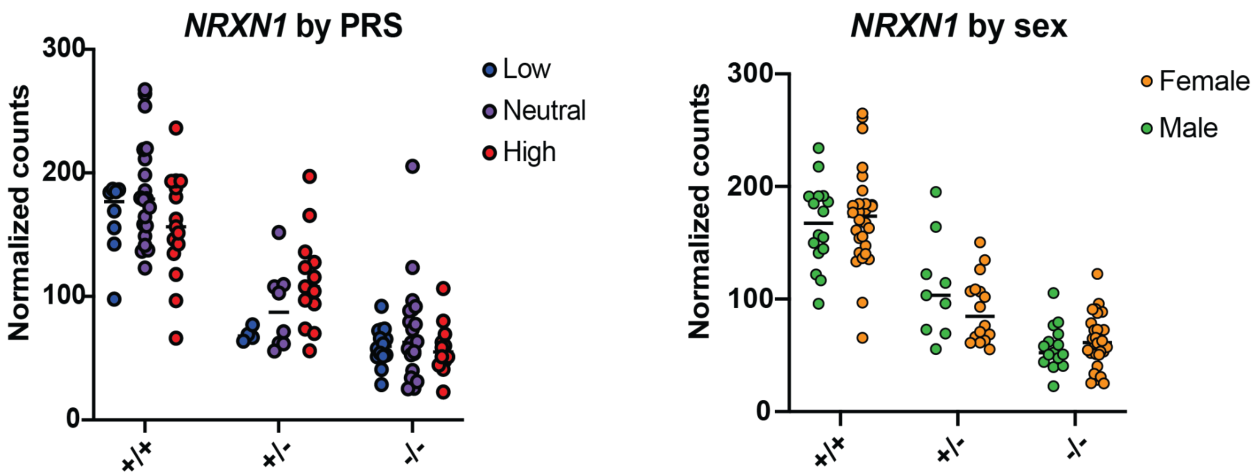


**Supplementary Fig. 4 | Examination of *NRXN1* expression by PRS and sex, related to Fig. 3.** Dot plots show *NRXN1* expression stratified by the PRS group (Left: blue = low, purple = neutral, red= high) and sex (Right: orange = female, green = male). A one-way ANOVA test was performed followed by Tukey's multiple comparison testing. There were no significant differences between PRS or sexes within genotypes.
