## Supplemental Figure 5 for "Gene editing in “cell villages” enables exploring disease-relevant mutations in many genetic backgrounds"

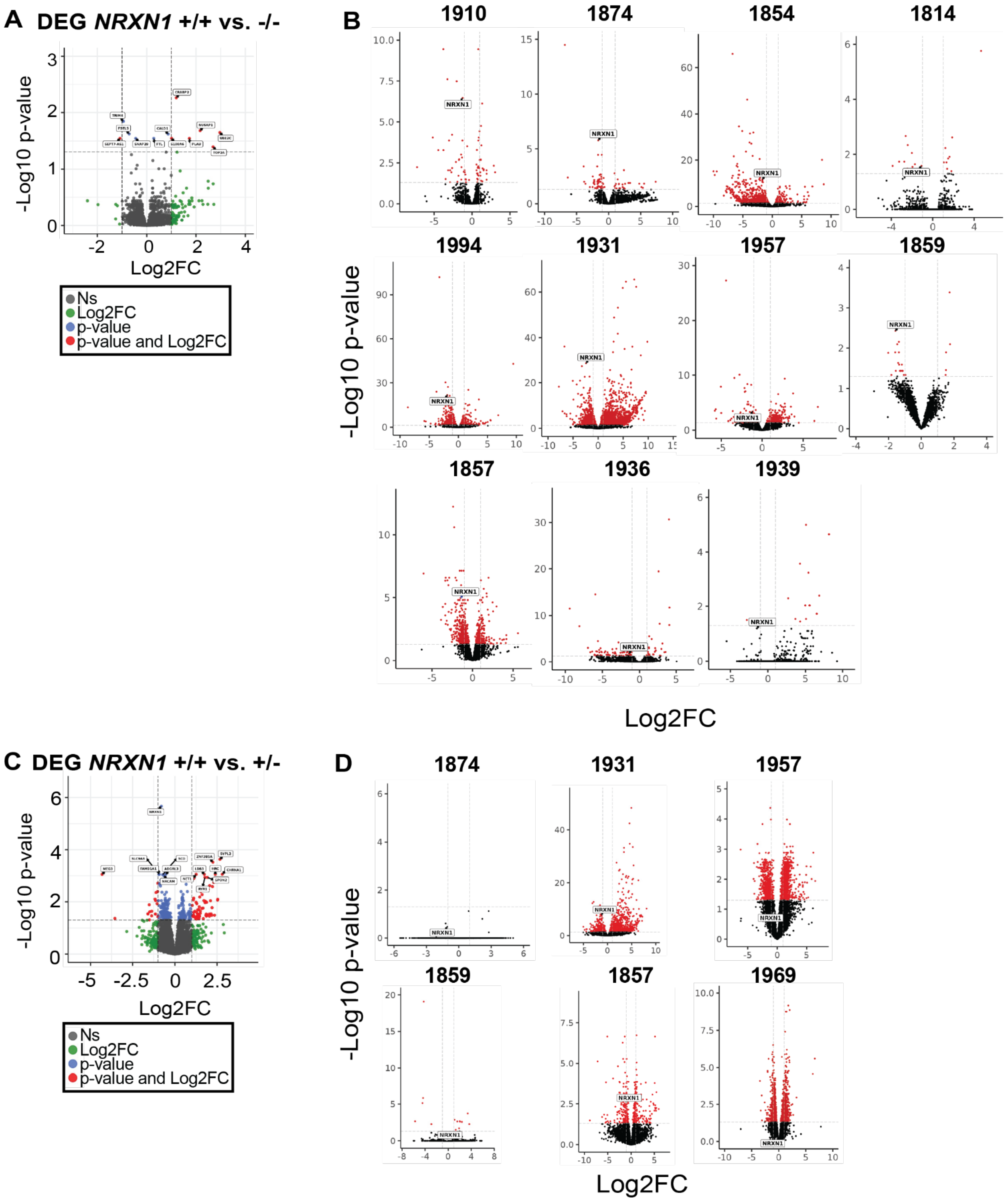


**Supplementary Fig. 5 | Differentially Gene Expression analysis of *NRXN1* -/- and +/- KOs compared to *NRXN1* +/+ neurons, related to Fig. 3. A, B** Volcano plots show differentially expressed genes in *NRXN1* -/- KO compared to unedited neurons analyzed together as a group (**A**) or individual (**B**). **C, D** Volcano plots show differentially expressed genes in *NRXN1* +/- KO compared to *NRXN1* +/+ neurons analyzed together as a group

(**C**) or individuals (**D**). The gene-level p-values were calculated using the Wald statistical test and adjusted for multiple testing using the Benjamini–Hochberg correction to control the false discovery rate (FDR) at 5%.
