## Supplemental Figure 6 for "Gene editing in “cell villages” enables exploring disease-relevant mutations in many genetic backgrounds"

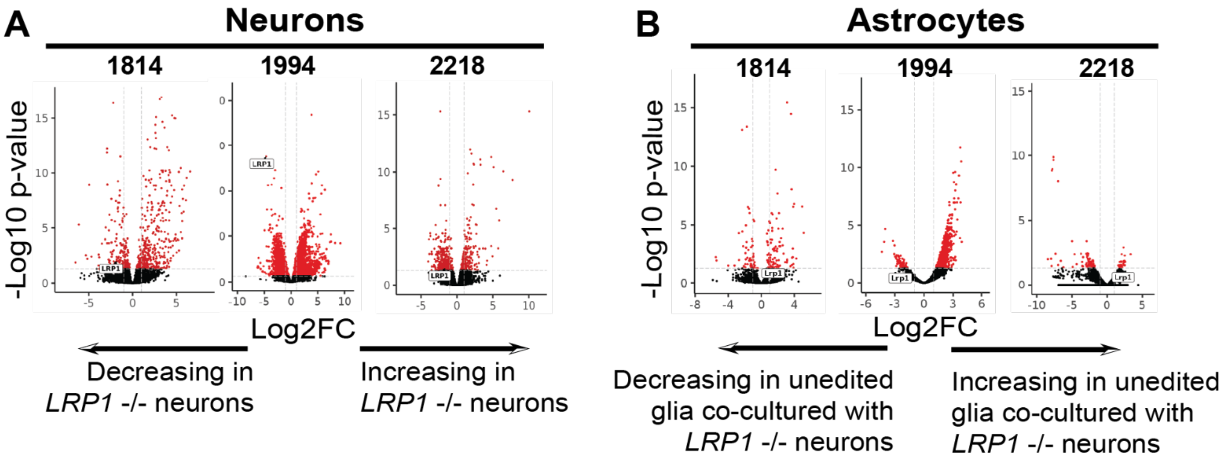


**Supplementary Fig. 6 | DEG analysis of *LRP1* -/- compared to *LRP1* +/+ neurons and co-cultured astrocytes, related to Fig. 4. A, B** Volcano plots show differentially expressed genes in *LRP1* -/- compared to unedited neurons (**A**) or co-cultured glia (**B**) analyzed by individual cell lines. The gene-level p-values were calculated using the Wald statistical test and adjusted for multiple testing using the Benjamini–Hochberg correction to control the false discovery rate (FDR) at 5%.
