## Supplemental Figure 7 for "Gene editing in “cell villages” enables exploring disease-relevant mutations in many genetic backgrounds"

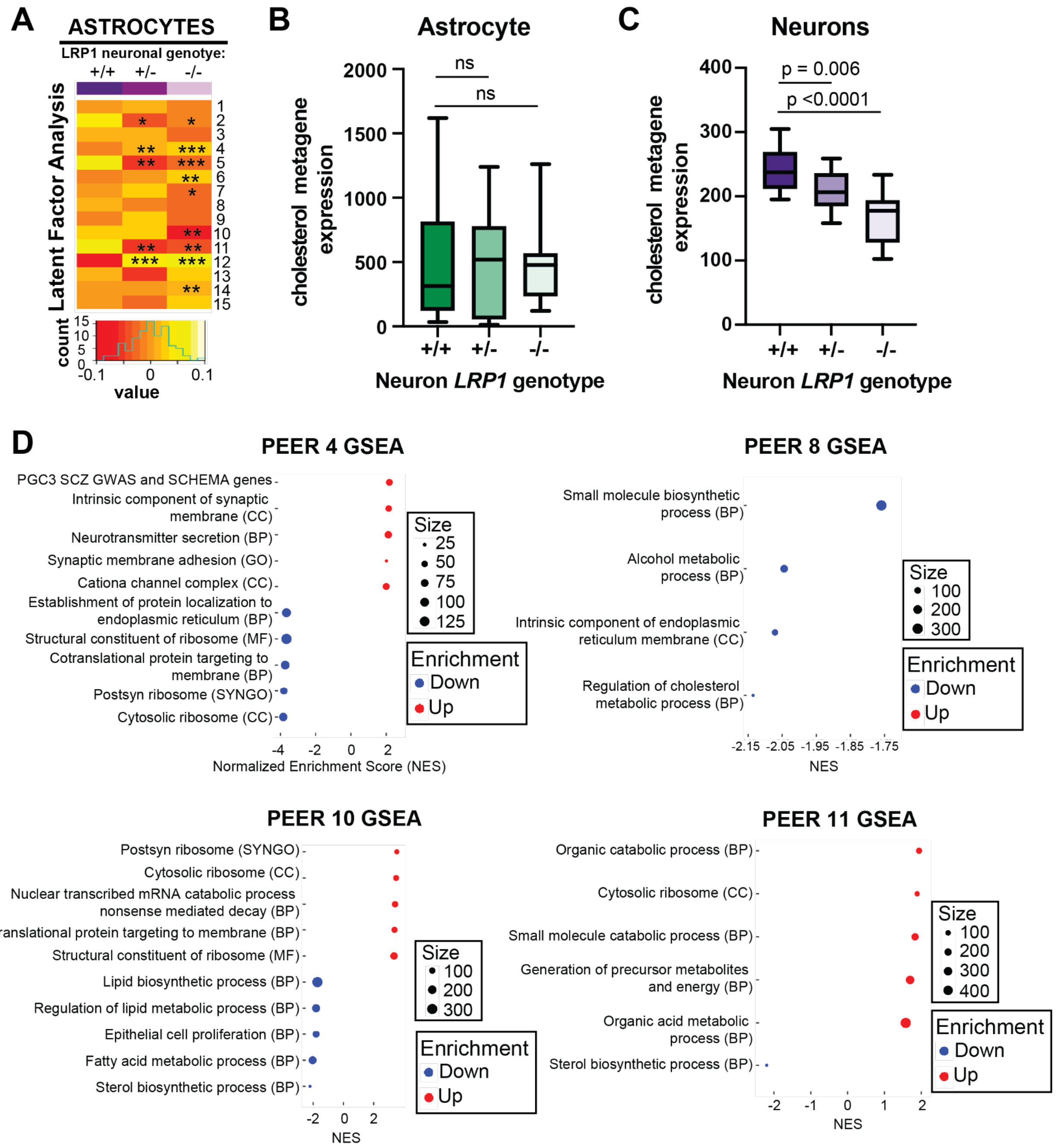


**Supplementary Fig. 7 | Latent factor analysis of mouse glia co-cultured with *LRP1* KO neurons, related to Fig. 4. A** Heat map displays latent factor counts in mouse glia co-cultured with neurons harboring the indicated *LRP1* genotype. **B, C** Cholesterol metagene expression in glia (**B**) and neurons (**C**). *$p < 0.05$, **$p < 0.01$, and ***$p < 0.001$ by Wilcoxon rank-sum test for panels A. The p values in B and C were calculated by one-way ANOVA followed by Tukey's multiple comparison test. **D** Gene set enrichment analysis for LF 4, 8, 10, and 11.
