## Supplemental Table 1 for "Gene editing in “cell villages” enables exploring disease-relevant mutations in many genetic backgrounds"

### Supplemental Table 1: Donor Information

| SCBB | Donor | Diagnosis | Sex | Ancestry | Ethnicity | PRS classification | Used in NRXN1 village | Used in LRP1 village |
|---|---|---|---|---|---|---|---|---|
| 1969 | CW20012 | Autism Spectrum Disorder | Female | White | Not Hispanic or Latino | high | yes | no |
| 2218 | CW20102 | Autism Spectrum Disorder | Male | White | Not Hispanic or Latino | high | yes | yes |
| 1814 | CW20103 | Control - Neurological - Family - Twin | Female | White | Not Hispanic or Latino | neutral | yes | yes |
| 1356 | CW20324 | Control - Neurological - Family | Male | White | Not Hispanic or Latino | neutral | yes | yes |
| 1910 | CW30154 | Heart Disease - Dilated Cardiomyopathy | Female | White | Not Hispanic or Latino | low | yes | yes |
| 1929 | CW30274 | Heart Disease - Dilated Cardiomyopathy | Male | White | Not Hispanic or Latino | high | yes | yes |
| 1994 | CW50037 | Control: No Cognitive Decline \| Non-Diabetic W | Female | White | Not Hispanic or Latino | neutral | yes | yes |
| 1605 | CW60221 | Intellectual Disability | Female | White | Not Hispanic or Latino | low | yes | yes |
| 1906 | CW70179 | Eye Disease - Age-related Macular Degeneratio | Male | White | Not Hispanic or Latino | low | yes | yes |
| 1859 | ML611-2911 | Control - Parent | Male | White | Not Hispanic or Latino | neutral | yes | yes |
| 1857 | ML611-3363 | Schizophrenia \| Schizoaffective \| Attention-Defic | Male | White | Not Hispanic or Latino | high | yes | no |
| 1854 | ML832-6778 | Control - Neurological - Parent | Female | White | Not Hispanic or Latino | low | yes | yes |
| 1864 | ML898-3533 | Control - Neurological - Parent | Male | White | Not Hispanic or Latino | low | yes | yes |
| 1874 | ML898-4425 | Control - Neurological - Parent | Female | White | Not Hispanic or Latino | low | yes | yes |
| 1867 | ML904-7021 | Schizoaffective | Male | Other/More than | Hispanic or Latino | high | no | yes |
| 1931 | ML907-2836 | Control - Neurological - Parent | Female | White | Not Hispanic or Latino | neutral | yes | yes |
| 1936 | ML909-6344 | Control - Neurological - Parent | Male | White | Not Hispanic or Latino | high | yes | yes |
| 1939 | ML910-8360 | Control - Neurological - Parent | Male | White | Not Hispanic or Latino | high | yes | yes |
| 1957 | ML911-9779 | Control - Neurological - Parent | Female | White | Not Hispanic or Latino | neutral | yes | yes |
