## Supplemental Table 2 for "Gene editing in “cell villages” enables exploring disease-relevant mutations in many genetic backgrounds"

### Supplemental Table 2: Primers and gRNAs

| Name | Gene targeted | Orientation | Sequence (5'→3') | Purpose | Efficiency | Selected for experiment |
|---|---|---|---|---|---|---|
| LD1166NRXN1s1 | *NRXN1* | sense | TTGATGTGTGCTGCTGTTGC | Sanger sequencing | N/A | N/A |
| LD1167NRXN1as1 | *NRXN1* | antisense | GTTTCTGGGGGAAGGAAGCA | Sanger sequencing | N/A | N/A |
| LD1168NRXN1s2 | *NRXN1* | sense | ACTGGGGCTTGTGGCTATTC | Sanger sequencing | N/A | N/A |
| LD1169NRXN1as2 | *NRXN1* | antisense | TACAACCTCCTGCTTCCAGC | Sanger sequencing | N/A | N/A |
| CB1308LRP1s | *LRP1* | sense | GTAAGGATGGACATGTTG | Sanger sequencing | N/A | N/A |
| CB1309LRP1s | *LRP1* | sense | TGATGAGTGCTCAGTGTAC | Sanger sequencing | N/A | N/A |
| CB1310LRP1as | *LRP1* | antisense | CCTCTGTTCAAGATAGGG | Sanger sequencing | N/A | N/A |
| hsGAPDH_F | *GAPDH* | sense | GTCTCCTCTGACTTCAACAGCG | qPCR | N/A | N/A |
| hsGAPDH_R | GAPDH | antisense | ACCACCCTGTTGCTGTAGCCAA | qPCR | N/A | N/A |
| hsRPLP0_F | RPL0 | sense | TGCCTCATATCCGGGGGAAT | qPCR | N/A | N/A |
| hsRPLP0_R | RPL0 | antisense | AAAAGGAGGTCTTCTCGGGC | qPCR | N/A | N/A |
| hsNRXN1_BF | *NRXN1* | sense | GAAGCCGTATTGGTGCGAGT | qPCR | N/A | N/A |
| hsNRXN1_BR | *NRXN1* | antisense | CGATGTCATCTGTCCCAACATT | qPCR | N/A | N/A |
| hsLRP1_AF | *LRP1* | sense | CTATCGACGCCCCTAAGACTT | qPCR | N/A | N/A |
| hsLRP1_AR | *LRP1* | antisense | CATCGCTGGGCCTTACTCT | qPCR | N/A | N/A |
| gRNA 1164 | *NRXN1* | N/A | GTACTGGGTCGGTCATTAGG AGG | gRNA | 900V: 15.2%; 1000V: 27.8%, 1100V: 44.3% | no |
| gRNA1165 | *NRXN1* | N/A | ACTGTCCACTCGCACCAATA CGG | gRNA | 900V: 31.4%; 1000V: 60.6%, 1100V: 89.4% | yes |
| gRNA1306 | *LRP1* | N/A | ATCTTGGCCACGTACCTGAG TGG | gRNA | 900V: 0%; 1100V: 18%, 1300V: 50% | yes |
| gRNA1307 | *LRP1* | N/A | GAGTTGGCTATCAACAGCAC AGG | gRNA | 900V: 0%; 1100V: 23%, 1300V: 60% | no |
