## Supplemental Table 3 for "Gene editing in “cell villages” enables exploring disease-relevant mutations in many genetic backgrounds"

**Supplemental Table 3: Summary of gene editing efficiency in cell village**

| Genotype/SCBB | Plate 1 | Plate 2 | Plate 3 | Plate 4 | Total |
|---|---|---|---|---|---|
| *NRXN1 +/+* | 47 | 1 | 3 | 1 | 52 |
| *NRXN1 +/-* | 13 | 3 | 1 | 1 | 18 |
| *NRXN1 -/-* | 10 | 34 | 27 | 38 | 109 |
| *NRXN1 fail* | 4 | 20 | 41 | 25 | 90 |
| *NRXN1 other* | 22 | 38 | 24 | 31 | 115 |
| *LRP1 +/+* | 23 | 16 | 21 | NA | 60 |
| *LRP1 +/-* | 13 | 17 | 17 | NA | 47 |
| *LRP1 -/-* | 33 | 33 | 29 | NA | 95 |
| *LRP1 fail* | 6 | 3 | 3 | NA | 12 |
| *LRP1 other* | 21 | 27 | 26 | NA | 74 |
